# VO: The Vaccine Ontology

**DOI:** 10.1101/2025.08.12.669998

**Authors:** Jie Zheng, Asiyah Yu Lin, Anthony Huffman, Anna Maria Masci, Rebecca Racz, Guanming Wu, Kallan Roan, Edison Ong, Sirarat Sarntivijai, Joy Hu, Eliyas Asfaw, Hayleigh Kahn, Xingxian Li, Xumeng Zhang, Nilufer Kosar, Jianfu Li, Warren Manuel, Rashmie Abeysinghe, Hasin Rehana, Benu Bansal, Yuanyi Pan, Jinjing Guo, Virginia He, Justin Song, Andrey I. Seleznev, Katelyn Hur, Anna He, Alexander Davydov, Qi Yang, Randi Vita, Bjoern Peters, Alan Ruttenberg, Alexander D. Diehl, Charles Tapley Hoyt, Paola Roncaglia, Rachael P. Huntley, Richard H. Scheuermann, Melanie Courtot, Thomas Todd, Samantha Sayers, Fang Chen, Xinna Li, Feng-Yu Yeh, Zuoshuang Xiang, Arzucan Ozgur, Patricia L. Whetzel, Mark A. Musen, Christopher J. Mungall, Wolfgang W. Leitner, Licong Cui, Lesley A. Colby, Harry L.T. Mobley, Brian D. Athey, Gilbert S. Omenn, Lindsay G. Cowell, Cui Tao, Junguk Hur, Barry Smith, Yongqun He

## Abstract

With the widespread use of vaccines in research and clinical settings, there is an urgent need to standardize vaccine representation, integrate information across diverse vaccine types, and support computer-assisted reasoning. Accordingly, we have since 2007 developed the community-based Vaccine Ontology (VO), which aligns with the Basic Formal Ontology and adheres to OBO Foundry principles. VO models ontologically vaccines, vaccine components, vaccine immune responses, vaccine investigation studies and other vaccine-related topics. VO represents more than 10,000 vaccines targeting 289 infectious pathogens and cancers in humans and over 30 nonhuman animal species. VO provides mappings to external resources such as RxNorm, CVX, FDA, and USDA. Various VO use cases exist. VO facilitates vaccine standardization in resources such as the VIOLIN vaccine database, ImmPort, and the Vaccine Adjuvant Compendium (VAC). Semantic queries can be made to query VO. VO has been shown to enhance experimental and clinical vaccine data analysis and vaccine literature mining. Overall, VO standardizes vaccine modeling and representation and greatly supports vaccine AI research in the Semantic Web era.

## Introduction

The invention and use of vaccines have dramatically improved human and other animal health over the past two centuries. There have been significant achievements in public health based on the population-wide administration of vaccines for smallpox, polio, measles, mumps, Rubella, diphtheria, tetanus, pertussis, influenza, and, more recently, varicella zoster (chickenpox and shingles) and hepatitis B, among others^1^. The impact of vaccines on the prevalence of formerly common childhood diseases is unparalleled by any other medical intervention in the history of humanity, with only the availability of clean water and sanitation having a similar effect^2^. Recently, vaccines have played a crucial role in combating the COVID-19 pandemic, reducing global deaths by over 60% during the first year of COVID-19 vaccination^3^. The widespread administration of these vaccines has significantly mitigated the impact of the pandemic and set a precedent for the rapid vaccine development in response to future global health threats.

Many vaccines have been approved and licensed for human or other animal use and have been recorded in various resources. The Centers for Disease Control and Prevention (CDC) developed and maintains the Codes for Vaccine Administered (CVX), a system in which each vaccine is given a specific code^4^. The U.S. Food and Drug Administration (FDA) has approved over 100 vaccines for human use^5^, each linked to a unique submission tracking number. The National Library of Medicine has developed the RxNorm vocabulary^6^, which standardizes the naming of prescription medications approved for human use in the United States. RxNorm includes thousands of human vaccines used in the U.S. For the standardization of vaccines not approved in the U.S., the Observational Health Data Sciences and Informatics (OHDSI) initiative developed the RxNorm Extension (RxE)^7^, which includes drug and vaccine terms used outside the U.S. and not covered by RxNorm. OHDSI harmonizes health data using the Observational Medical Outcomes Partnership (OMOP) Common Data Model (CDM) and its standardized vocabularies^7,8^. The U.S. Department of Agriculture (USDA) has approved about 1000 veterinary vaccines for use within the country, assigning each a unique code^9^. But where these and other organizations have assigned codes or identifiers for easy reference and data exchange within a specific domain or context, these standards are inherently not interoperable, preventing advanced seamless data integration and analysis. This is the problem that the VO is designed to address.

Beyond already approved vaccines, thousands more, especially cancer vaccines, are in various stages of clinical trials or pre-trial laboratory research. Data about these trials can be found on resources like clinicaltrials.gov^10^ and the NCI Thesaurus^11^. Furthermore, extensive research has been conducted on vaccines and vaccination. Currently, over 550,000 vaccine-related articles are available in PubMed^12^, reflecting the depth and breadth of ongoing inquiry in this critical area. With the large amount of vaccine data available in various resources, developing an efficient strategy for vaccine data retrieval, standardization, and integration is challenging.

Biomedical ontologies are sets of terms and relations that represent entities in the biomedical world and how they relate to each other. Ontology terms form a tree, representing classes of entities at different levels of generality. Each term is associated with documentation, including alphanumeric VO identifiers, labels, and definitions, which are expressed both in human-readable form and in a logical language to enable support for automated reasoning. In the past two decades, hundreds of biomedical ontologies, such as the Gene Ontology (GO)^13,14^ and the Ontology for Biomedical Investigations (OBI)^15,16^, have been developed to provide controlled and standardized terms for used in different biological and biomedical domains^17,18^. These terms are then used to annotate in consistent fashion hundreds of different types of data, whereby the resultant annotation-based data repositories for a highly valuable resource for both querying and analyzing high-throughput data. Biomedical ontologies have been widely used in multiple applications, including data harmonization, integration, and computational data analysis and reasoning, and they have also advanced artificial intelligence (AI) and machine learning (ML) research based on big biological and biomedical data. For example, a deep neural network model predicts protein function from sequence by leveraging both GO annotations and the hierarchical structure of the GO^19^. The Human Disease Ontology (DOID)^20–22^ has been applied to improve the accuracy of Named Entity Recognition (NER) systems based on deep learning^23^.

To promote vaccine data standardization, integration, analysis, and computer-assisted reasoning, we have developed the Vaccine Ontology (VO) in a collaborative, community-based fashion following the Open Biological and Biomedical Ontology (OBO) Foundry principles^24,25^. The development of the ontology began in 2007, led by a team of domain and ontology experts representing various groups whose work required standardized representations of vaccine information. The VO was first presented at the 1st International Conference on Biomedical Ontology in 2009 in Buffalo, NY, USA^26^. Over the past 18 years, more than 400 papers have cited VO^27^. However, this is the first journal article designed to introduce the vaccine ontology systematically in order to give users a comprehensive understanding of VO, its overall design, high-level terms, and hierarchy, and to provide examples of how VO has been used to support data standardization and to address research questions in the vaccine domain.

## Results

### High-level VO classes and their alignment with reference ontologies

Figure 1 illustrates the high-level hierarchical structure of the VO. Overall, all VO terms are aligned with the Basic Formal Ontology (BFO)^28^, a top-level ontology conformant to the ISO/IEC 21,838 standard^29^. As a domain-neutral framework, BFO includes two main branches: continuant and occurrent (Figure 1). A continuant is an entity that persists, endures, or continues to exist through time while maintaining its identity. Independent continuants, such as vaccines and organisms, can exist by themselves, while dependent continuants, such as quality and function, depend for their existence on the independent continuants. Occurrents include time intervals and any entity that unfolds in a time-dependent manner, such as a process of vaccination or of response to a vaccine. More than 600 ontologies have adopted BFO^30^, and the adoption of BFO by VO thus supports its broad interoperability with ontologies in other domains.

**Figure 1.**
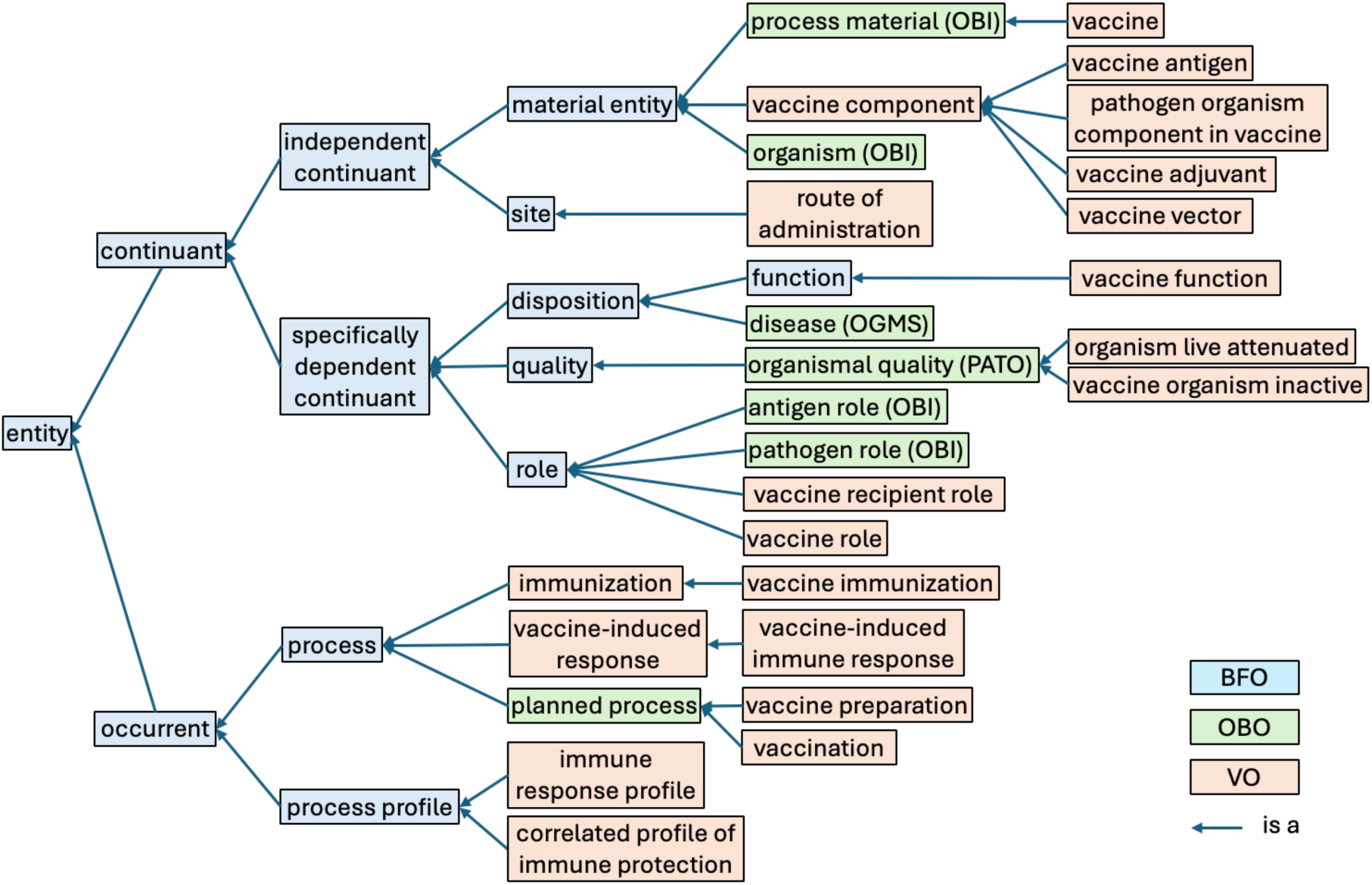
High-level structure of VO. (abridged: some intermediate non-VO terms omitted for simplicity)

VO reused many reference ontologies in the OBO library^17^, including the OBI^15,16^, the Phenotype and Trait Ontology (PATO)^31^, and the Ontology for General Medical Science (OGMS)^32^ (Figure 1). OBI provides a set of terms commonly used in biomedical investigations, for example, materials used in processes, organisms, antigen and pathogen roles, assay and analysis types^15,16,33^. PATO describes various phenotypes and qualities^31^, and OGMS defines general terms used in clinical encounters such as disease, sign, and symptom^32^.

New VO terms can be created in a semantically uniform way under the guidance of the VO high-level structure (Figure 1) and by the BFO and intermediate layers of OBO reference ontologies (e.g., OBI, PATO, and OGMS) incorporated therein. Under the BFO:continuant branch are: vaccines, vaccine components (e.g., vaccine antigen and adjuvants), sites (e.g., route of administration), and vaccine-associated functions and roles. Under the BFO:occurrent branch are: vaccine preparation, vaccination, vaccine immunization, vaccine-induced immune responses, and vaccine immune profiles, etc. The definitions and associated classifications of these terms are detailed below.

### Ontological modeling of vaccine hierarchies using VO

VO defines ‘vaccine’ (VO:0000001) as “material entity that is manufactured to realize the vaccine function.” The definition emphasizes the ‘vaccine function’ to distinguish ‘vaccine’ from other manufactured materials such as ‘devices’ that have functions such as containing, measuring, and so forth. In addition, ‘vaccine’ itself is a processed material that excludes those natural pathogen organisms that can induce immune responses.

Figure 2 illustrates the classification of various vaccine types and their hierarchies in the VO. The central branch of the hierarchy is labelled ‘vaccine against disease.’ Terms in this branch categorize vaccines based on the targeted disease under headings such as infectious disease, cancer, and allergy. Traditionally, vaccines against infectious diseases are classified based on the type of causative agent: bacteria, virus, or eukaryotic microorganism such as fungi and protozoans. To classify the associated vaccines, VO deploys two parallel definition strategies: 1. based on the infectious disease type (for example: tuberculosis, AIDS) and 2. based on the underlying causal pathogens (for example *Mycobacterium tuberculosis*, human immunodeficiency virus).

**Figure 2.**
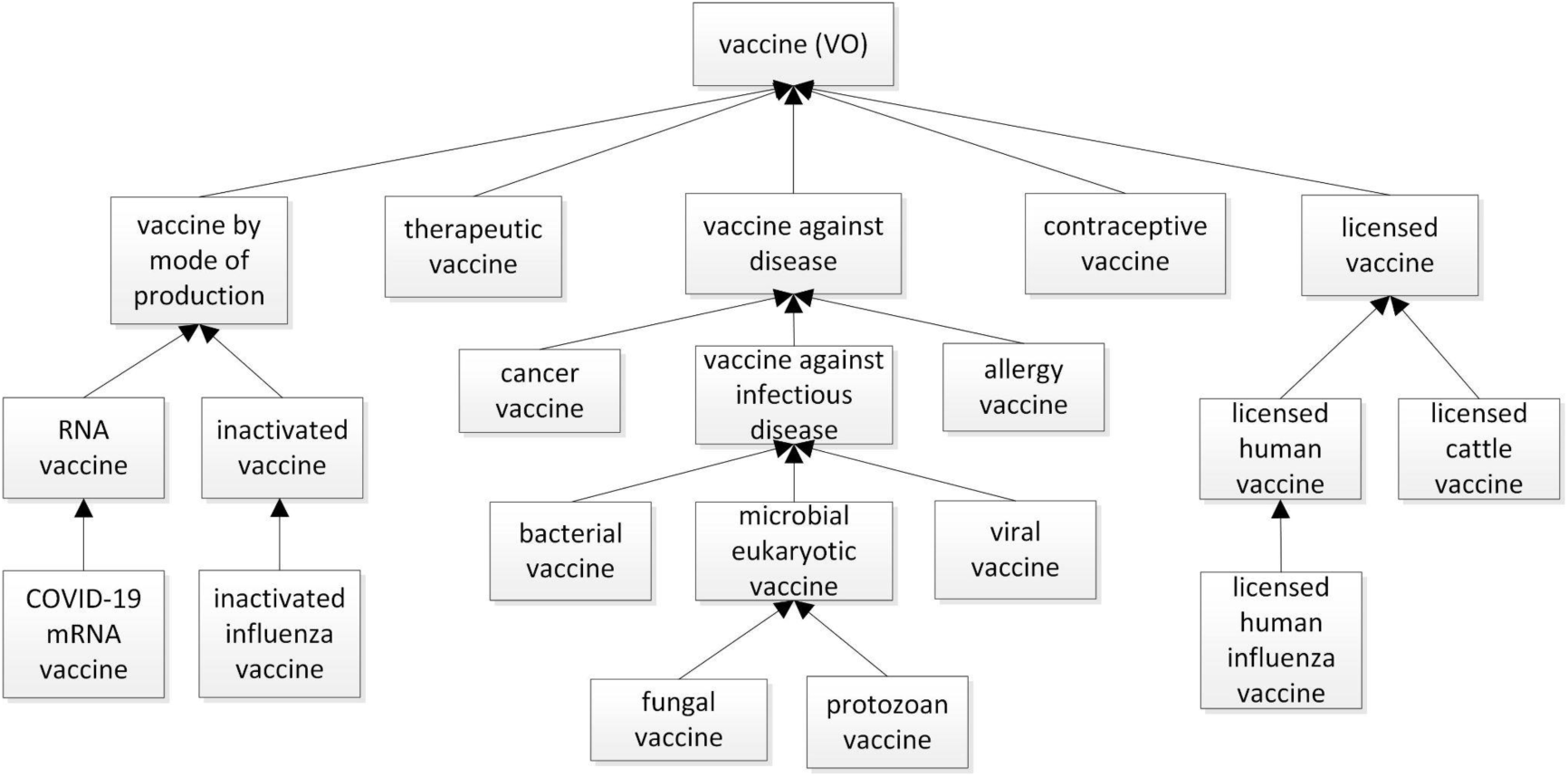
Representative vaccine types and their associated hierarchies as defined in the VO.

Different vaccine types are often classified based on their roles. Overall, the ‘vaccine role’ is a role that inheres in a prepared material entity that is designed to induce protection or treatment for a disease or infection. VO includes first vaccine roles such as the ‘licensed vaccine role’ and ‘authorized vaccine role,’ which are assigned to vaccines by specific organizations. Additionally, VO uses roles such as ‘vaccine in clinical trial role,’ and ‘vaccine in research role,’ alongside the licensed and authorized vaccine roles, to signify the life cycle phase of a vaccine (Figure 2).

Vaccines are often classified on the basis of the mode of vaccine production (Figure 2). This reflects the fact that vaccines can be prepared using different methods or technologies, which reflect how the vaccine delivers antigens to the recipient’s immune system. Different vaccine preparation methods produce vaccines with specific roles, which are acquired during the process of preparation and realized in the process called ‘vaccine immunization.’ These diverse vaccine roles facilitate logical inference by the ontology reasoner. Further examples include: ‘live attenuated vaccine role,’ ‘inactivated vaccine role,’ ‘subunit vaccine role,’ ‘RNA vaccine role,’ ‘DNA vaccine role,’ ‘conjugate vaccine role,’ ‘toxoid vaccine role,’ and ‘virus-like particle vaccine role.’ Each of these terms indicates a corresponding type of vaccine, such as ‘live attenuated vaccine,’ ‘inactivated vaccine,’ and so forth. VO defines the latter in terms of the former as a strategy to avoid the forking of data which arises when a single ontology term has two direct parents within the ontology hierarchy.

Vaccines can also be defined using other characteristics. For example, ‘therapeutic vaccine’ and ‘preventive vaccine’ are defined based on two types of vaccine functions, where the function of an entity is defined in terms of the entity’s reason for existence (the purpose it is designed to serve). While most existing vaccines are preventive, many therapeutic vaccines are being developed and tested in areas such as control against various cancer types^34,35^. Examples of other vaccine types are: contraceptive vaccines^36,37^, and editable vaccines, which are produced in genetically modified plants or other foods, which can be eaten to deliver immunization against specific diseases^38,39^. Prime-boost vaccines deploy a vaccination strategy that uses different vaccine types for the initial (prime) and subsequent (boost) doses to enhance the immune response by combining the strengths of multiple vaccine platforms^40,41^.

### Ontological modeling of vaccine components and formulations using VO

A vaccine typically contains multiple ingredients; each defined as a ‘vaccine component’ in the VO hierarchy (Figure 1). The most critical component of a vaccine is either an antigen, or it is some other molecule (e.g., RNA or DNA) that can be transcribed and translated into an antigen, which is recognized by receptors of the immune system to induce an immune response. Other vaccine components identified in VO include: ‘pathogen organism component in vaccine,’ ‘vaccine vector,’ ‘vaccine adjuvant,’ ‘vaccine stabilizer,’ and ‘vaccine preservative.’ These terms have been described in detail in our previous publication^42^. The vaccine formulation method, the process of combining various components to produce an administrable vaccine, is closely associated with the provision of profiles of vaccine safety and efficacy^42,43^.

The latest (2025-07-06) VO release comprises some 230 vaccine adjuvants, representing a major advance over previous versions. Sources of these new adjuvants include the Vaccine Adjuvant Compendium (VAC)^44^ and direct literature annotations. Vaccine adjuvants have been classified hitherto primarily on the basis of their sources and of their chemical and physical characteristics. VO captures in addition, again by using logically formulated axioms, the associated induced immune profile properties. A given vaccine adjuvant can induce either one or multiple distinct immune profiles. For example, ‘LTA1 vaccine adjuvant’ induces three different ‘immune profiles’: Th1-biased, Th2-biased, and Th17 immune profile^45^ (Figure 3a). A Description Logic (DL) query is a formal expression used to retrieve or infer information about entities within an ontology based on logical constructs in the formal ontology^46^. DL queries based on these three different ‘immune profiles’ revealed that the ‘LTA1 vaccine adjuvant’ is associated with all three immune profile types (Figure 3b), demonstrating the use of DL queries for automated knowledge discovery. Note that DL queries and results shown in Figure 3b are available at the Zenodo repository^47^. In future VO releases we will represent additional vaccine adjuvant properties, such as receptors and routes of immunization, to enable the identification of vaccine adjuvants based on these characteristics.

**Figure 3.**
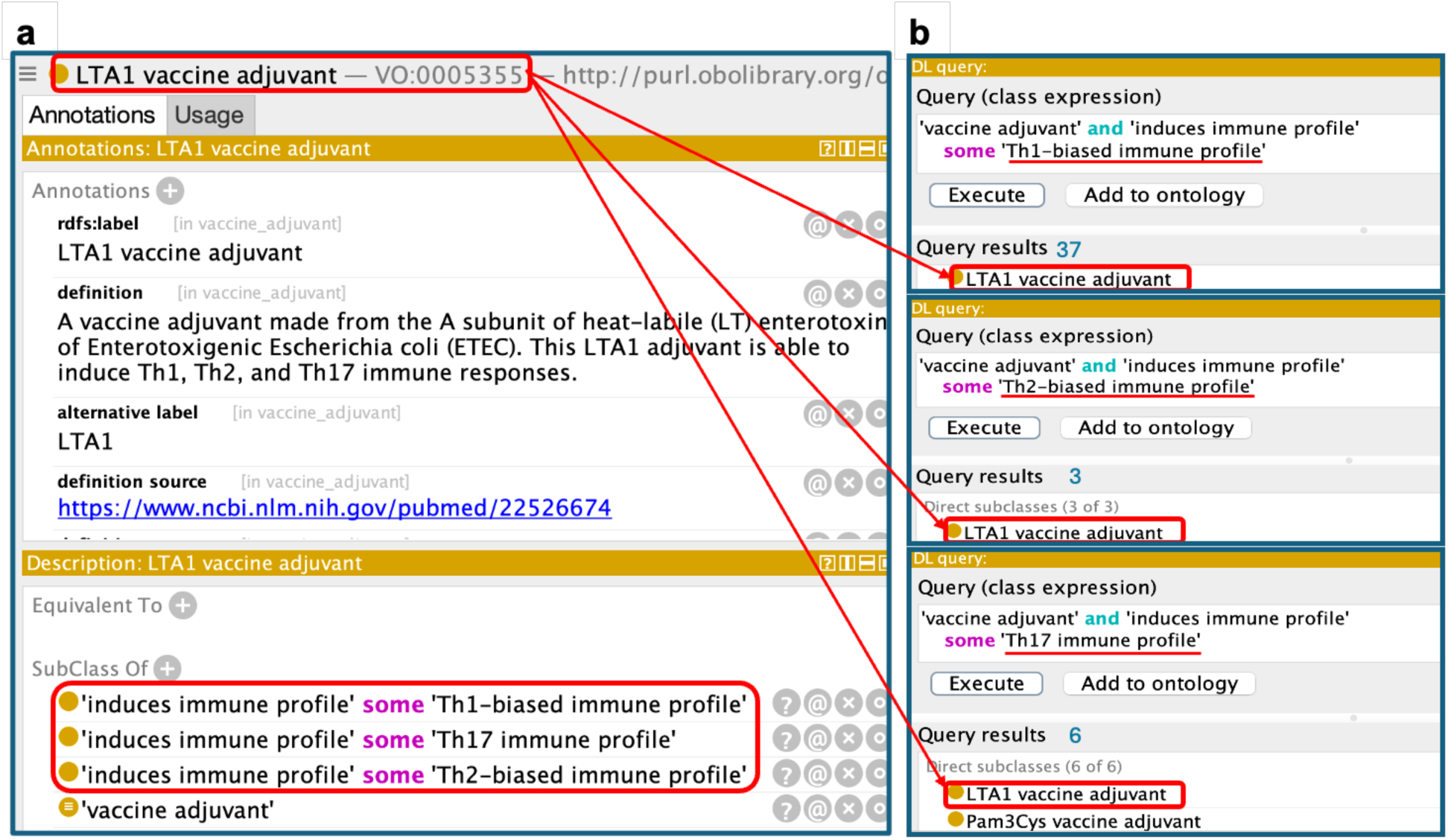
VO representation of the LTA1 vaccine adjuvant. (**a**) VO representation of the adjuvant, including three immune profiles induced by the adjuvant. (**b**) DL queries constructed from these three ‘immune profiles’ successfully retrieve the LTA1 adjuvant.

### Ontological modeling of individual vaccines based on the VO ontology design pattern

An ontology design pattern (ODP) is a reusable template used to define a specific type of entity^48,49^. Figure 4a illustrates the vaccine design pattern that covers features shared by most vaccines. Specifically, a vaccine can be defined or categorized^50^ along a number of dimensions, including: 1) vaccine immunized recipient, 2) vaccine targeted disease or pathogen, 3) vaccine function, 4) vaccine components, 5) vaccine development status, 6) vaccine manufacturer, and 7) vaccination routes. As the output of the ‘vaccine preparation’ process, a ‘vaccine’ contains multiple components, such as a protein component with ‘vaccine antigen role’ and a ‘vaccine adjuvant.’ Roles, functions, and process profiles are often used for vaccine knowledge representation. For example, a ‘vaccine’ is administered to the organism that has the ‘vaccine recipient role’ at a ‘route of administration’ site. After the ‘vaccination’ process there follows a ‘vaccine immunization’ that is induced by the vaccination of the ‘vaccine,’ and this in turn has outcome ‘vaccine-induced host immune response’ with a process profile ‘correlated profile of immune protection.’ The ‘vaccine function’ is realized through ‘vaccine immunization’ immunization against ‘pathogen’ infection and disease. In VO, the vaccine manufacturers and vaccination route are generally indicated in connection with a licensed vaccine product (Figure 4a).

**Figure 4.**
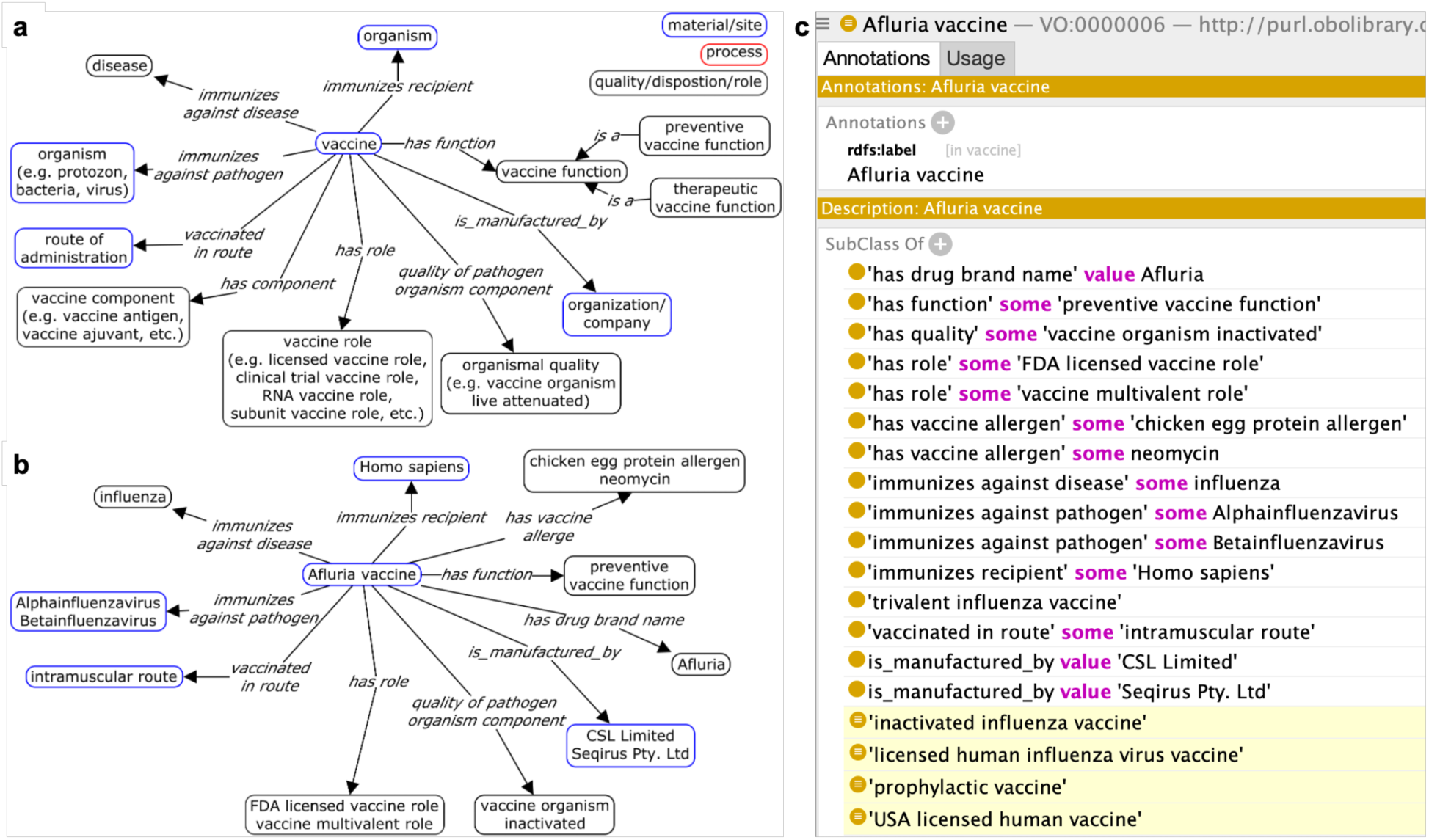
Vaccine ontology design pattern (ODP) in VO. **(a)** ODP of a typical vaccine. **(b)** Illustration of vaccine ODP with ‘Afluria vaccine.’ **(c)** Representation of ‘Afluria vaccine’ in VO with inferred parents highlighted in yellow.

Our vaccine ODP effectively addresses the complexity of multiple inheritance by asserting a vaccine under a single parent, namely that defined by the vaccine-targeted diseases or pathogens, and utilizing an OWL reasoner to infer additional parents. A specific vaccine can be classified simultaneously via multiple of the above-listed categories. For example, a vaccine can be simultaneously an ‘influenza virus vaccine,’ ‘inactivated vaccine,’ and ‘licensed human vaccine.’ Multiple inheritance occurs when a single vaccine is associated with multiple parent categories. To enhance clarity, modularity, and maintainability, it is greatly recommended to avoid asserting multiple parents in ontology development^51^. In VO, each vaccine is asserted under only one parent, with the vaccine-targeted diseases or pathogens being the major classification category. Remaining parents are automatically inferred based on an OWL reasoner such as ELK^52^.

Figures 4b and 4c illustrate how the vaccine ODP represents the ‘Afluria vaccine.’ First, the term ‘Afluria vaccine’ is asserted under ‘trivalent influenza vaccine,’ which is a subtype of influenza vaccine. The Afluria vaccine antigen is the whole viral organism that has the quality ‘inactivated’. The vaccine itself is asserted under a single parent in VO. We rely on OWL reasoners to infer additional parents. Based on the logical axioms governing ‘Afluria vaccine,’ a reasoner will classify it as an ‘inactivated influenza vaccine,’ ‘licensed human influenza virus vaccine,’ ‘prophylactic vaccine,’ and ‘USA licensed human vaccine.’

### Ontological modeling of vaccination and vaccine-induced immune responses in VO

In VO, the term ‘vaccination’ (VO:0000002) is defined as “Process of administering a vaccine *in vivo* to a recipient (e.g., a human), with the intent to invoke a protective or therapeutic adaptive immune response.” This definition features three OBI terms: ‘vaccination’, defined as a process of ‘administering substance in vivo’ (OBI:0600007), in which some ‘vaccine’ realizes the ‘material to be added role’ (OBI:0000319) to an organism (OBI:0100026).

Although often commonly considered as a synonym of ‘vaccination’, ‘immunization’ is defined with a distinct meaning. Specifically, VO defines ‘immunization’ (VO:0000490) as: “Process that results in an adaptive immune response to one or more antigens.” ‘Vaccine immunization’ refers to an immunization induced by a vaccine through the vaccination process.

VO defines various vaccine-related immune responses and immune profiles. ‘Vaccine-induced response’ is a process that specifies the organism’s response to an administered vaccine, which can occur at multiple levels: population, organism, system, cell, and gene/protein. The ‘vaccine-induced immune response’ is a specific type of this response. The ‘immune profile,’ a subclass of the BFO ‘process profile,’ represents a pattern within the immune response process, encompassing the interconnected parts and operations of the immune system that contribute to the overall immune reaction. Similarly, the ‘correlated profile of immune protection’ is another BFO ‘process profile’ that denotes measurable biological responses statistically linked to protection against specific infections or diseases. Both ‘immune profile’ and ‘correlated profile of immune protection’ are pivotal for researchers to comprehend the mechanisms of vaccine-induced immunity. More details about these are reported in our manuscript (under review)^53^.

### Ontological modeling of experimental vaccine investigation studies

Another major topic in VO is the ontological modeling of experimental vaccine studies. Two main types of such studies exist: vaccine immune response studies and challenge experiments for vaccine efficacy studies. To study the immune responses induced by a vaccine, groups of organisms (e.g., mice or humans) are administered the vaccine or a control agent (e.g., saline), followed by samples collected and assayed for specific immune responses^54^. The gold standard for measuring the efficacy of a preventive vaccine or vaccine candidate against an infectious agent is to determine whether it induces protection against a challenge infection with a virulent pathogen or its virulence factor^15,55,56^. The vaccine protection (or vaccine challenge) experiment involves three steps: (1) Vaccination, (2) Pathogen challenge, and (3) Assessment of vaccine protection outcome^15^. VO will advance ontological modeling of these studies, which will in turn enhance study understanding, facilitate the integration of data from multiple sources, identify variables that may affect study outcomes, and support statistical data analysis.

An example is the use of VO and the Ontology for Biological and Clinical Statistics (OBCS)^57^, to represent a general model of vaccine immune response studies and associated data analysis^54^. This general ontological model was applied to yellow fever vaccine immune response studies that employed transcription profiling assays by array to examine immune responses. The study’s metadata types were standardized using the mentioned ontologies to compare differences in similar studies^54^. Further analyses investigated why similar studies generated different gene expression results when induced by the same yellow fever vaccine^58^. Investigations of this sort are important for an improved understanding of vaccine-induced protective immunity.

The VO was utilized to represent live attenuated *Brucella* vaccine challenge studies, focusing on variables that may contribute to vaccine efficacy^59^. In total 74 peer-reviewed publications detailing the efficacy of live attenuated *Brucella* vaccines were manually curated based on the VO model. The variables and their values were extracted for ANOVA analysis, which identified nine variables that significantly contributed to the efficacy of *Brucella* vaccine protection^59^.

The Vaccine Investigation Ontology (VIO) is a subset of VO with the focus on ontological modeling of vaccine investigations^58^. VIO has been used to identify and to represent ontologically multiple variables used in vaccine investigation studies. Our VIO modeling identified different settings of variables in two gene expression profiling studies, authored by Gaucher *et al*.^60^ and Querec *et al*.^61^, which used samples collected from human subjects vaccinated with the same Yellow Fever vaccine 17D (YF-17D). VIO-based gene expression data analysis revealed that the significantly expressed gene lists from the two studies differed substantially, but that more uniform results were found when the Gene Ontology and pathway analyses were performed^58^. This study showed how different values for specific variables may explain different results obtained from similar studies.

### VO statistics

The latest release of the Vaccine Ontology (VO), version 2025-07-06^62^, contains a total of 15,383 terms. These include 14,509 classes, 122 object properties, 162 annotation properties, and 590 instances. Of these, 14,005 terms (91%) are specific to VO, including 13,345 classes, 46 object properties, 25 annotation properties, and 570 instances/individuals. The remaining terms are imported from external ontologies. Currently, VO has imported 1,144 classes from 22 external ontologies, such as the OBI^16,63^, NCBI organismal classification (NCBITaxon)^64^, DOID^20–22^, and the Ontology for Parasite Lifecycle (OPL) ^65^. Detailed statistics regarding VO terms based on their respective prefixes are available at the Zenodo repository^47^ (VO_statistics-v2025-07-06.xlsx).

Most VO terms designate vaccines. As of 2025-07-06^62^, VO represents 10,940 vaccines or vaccine candidates targeting either humans or one or more of 33 different genera or species of other animals. While most of these vaccines (9,983) are directed against 289 infectious pathogens, some target non-infectious diseases, including 677 cancer vaccines, 4 arthritis vaccines, 4 diabetes vaccines, and 2 autoimmune disease vaccines.

**Table 1** summarizes the distribution of vaccines according to their term resources and their development phases. 7,863 VO vaccines are licensed or authorized for use, 805 are vaccine candidates in the clinical trial stage, and 1,273 are vaccine candidates in the research phase. Both licensed human and veterinary vaccines were categorized. For licensed human vaccines, we have collected 122 FDA-approved vaccines (retrieved on February 18, 2025)^5^, 272 CVX codes (extracted on May 22, 2024)^4^, 2,051 vaccine-related terms from RxNorm (version 2023-07-03)^66^, and 5,389 vaccine-related terms from the RxE (version 2023-08-24)^66^. Currently, VO contains all FDA-approved vaccines, the complete CVX code set^67^, the full list of RxNorm terms (including vaccines and vaccine ingredients)^66^, and 4,091 terms from the RxE^66^. In addition to licensed human vaccines, VO also includes 790 USDA vaccines licensed for veterinary use. There are approximately 1,000 vaccines in the USDA catalog (released on January 30, 2025), and efforts are ongoing to incorporate into VO all licensed veterinary vaccines from the catalog. VO vaccine stages from research and clinical trials to licensed/authorized use are illustrated in **Table 1**.

**Table 1.** Summary of vaccines by development phase and term resource.

| Development phase | Target host | Term resource | Vaccine Count | Total |
| --- | --- | --- | --- | --- |
| Licensed/Authorized | Human | FDA | 115 | 7,054 |
|  |  | CVX code | 271 |  |
|  |  | RxNorm | 2,029 |  |
|  |  | RxNorm Extension | 4,064 |  |
|  |  | Other | 575 |  |
|  | Veterinary animals | USDA | 789 | 809 |
|  |  | Other | 20 |  |
| In Clinical trial | Human | clinicaltrials.gov & NCI Thesaurus | 427 | 805 |
|  | Human | Other | 378 |  |
| In Research | Human + Veterinary animals | Literature and VO users | 1,273 | 1,273 |

To automate VO development, we created 19 ROBOT templates based on the VO ODPs. ROBOT^68^ is a command-line tool used to automate ontology development workflows, especially for OBO Foundry ontologies. A ROBOT template can be used for standardized data annotation, and the standardized data can be processed by ROBOT for automatic ontology term generation, ontology integration, and format conversion. These 19 VO-specific ROBOT templates were converted to the corresponding modules using the ROBOT tool for integration into the VO. These modules encompass vaccine, vaccine adjuvant, non-adjuvant vaccine components, VO individuals, VO object properties, and VO annotation properties. Five templates/modules were designed to support the mapping of annotations between VO and external vaccine data resources. Detailed descriptions of the templates are available at: https://github.com/vaccineontology/VO/wiki/Editing-VO#vo-templates.

### VO records cross-referencing and mapping to external data resources

VO references terms collected from external terminologies or resources using different annotation properties. For example, VO uses the annotation properties ‘CVX code’ and ‘RxNorm ID’ to cross-reference CVX and RxNorm identifiers for specific vaccines. The related information from original sources of these terms, including their original unique identifiers and names or labels, has been collected in the ROBOT templates and incorporated into the VO. These mappings, which represent relationships between terms in different ontologies, vocabularies, or classification systems, are a significant contribution to data interoperability and integration across diverse vaccine domains and resources.

To further enhance vaccine data exchange and support text mining, we generated term mappings between VO and external resources using the Simple Standard for Sharing Ontological Mappings (SSSOM) standardized format^69^. The SSSOM format mapping files were developed from the ROBOT templates include: VO to RxNorm vaccines, VO to CVX codes, VO to concept IDs in OHDSI vocabularies (including RxNorm, RxE vaccines, and CVX codes), VO to FDA vaccines, VO to USDA veterinary vaccines, and VO to VAC vaccine adjuvants. These files are available on the GitHub site: https://github.com/vaccineontology/VO/tree/master/mappings.

### Use cases of the VO

The VO has been used in many applications. Here we illustrate how it has been applied in five specific areas:

***(1) Vaccine data standardization for various vaccine-related resources***

The initial source of VO development is the Vaccine Investigation and Online Information Network (VIOLIN), an online resource focused on vaccines and vaccine-related data^55,56^ pertaining to vaccine components (such as vectors and adjuvants), preparation methods, targeted pathogens and diseases, species for which they are licensed, and manufacturers. VO has been used to standardize the representation of vaccines, vaccine adjuvants, vaccine vectors, vaccine types based on production methods, and routes of vaccine administration drawing on the integrated data in VIOLIN covering the complex mechanisms underlying vaccine function^70–73^.

VO itself has been reused in many vaccine-related resources. For example, the Immunology Database and Analysis Portal (ImmPort) is an NIH-funded resource that aims to promote the sharing and exchange of both raw data and analyses within the immunology community^74^. The ImmPort database employs terminologies and ontologies to harmonize diverse immunology studies, ensuring consistency and comparability. VO is used to promote standardized vaccine and vaccine adjuvant annotation in ImmPort and significantly facilitates advanced vaccine immune response analysis (covered in more detail below in Section (3)). Examples of other resources using VO include the ImmuneSpace repository developed by the Human Immunology Project Consortium (HIPC)^75^, the Immune Epitope Database (IEDB)^76,77^, and the Vaccine Adjuvant Compendium (VAC)^44^. The usage of standardized VO terms in these resources enables data harmonization and enhances the FAIRness^78^ – Findability, Accessibility, Interoperability, and Reusability – of the databases in question.

The Vaccine Adjuvant Compendium (VAC) is an intramural program of the National Institute of Allergy and Infectious Diseases (NIAID), and the VAC team sought support from the VO development group upon its initiation in 2021. The VAC database catalogs novel, accessible vaccine adjuvants, including those developed with NIAID support. The database collects and displays the characteristics and metadata pertaining to vaccine adjuvant types in order to help vaccine developers identify adjuvants suitable for use in vaccines against given immune-mediated and infectious diseases and against cancer^44,79^. Data entered into VAC details properties of an adjuvant, such as its chemical nature, receptor, induced immune profile, and route of immunization, as well as associated publications and contact information for establishing agreements. VO’s developers have collaborated closely with the VAC team to systematically represent each VAC adjuvant record in VO. Each VAC record is represented in VO with a unique identifier, and the vaccine adjuvant-induced immune profile (e.g., Th1-biased immune profile) is also represented (Figure 3), supporting standardized annotation and query. Of 230 vaccine adjuvants in VO, 87 are sourced from the VAC.

***(2) Queries on the VO and VO-based vaccine knowledgebase***

As VO systematically represents vaccines, vaccine components, vaccine responses, and other related entities, the VO itself can serve as a semantic vaccine knowledgebase. Developed using the W3C standard Web Ontology Language (OWL)^80^, automated inferences from (“reasoning”) and analysis of vaccine data represented in VO. When the VO is opened in an OWL editor such as the Protege OWL editor^81^, we can use the semantic query method DL query^46,82^ to query the VO knowledge as demonstrated in Figure 3. When VO is stored in a triple store such as the Ontobee linked ontology repository and data server^83,84^, SPARQL (i.e., SPARQL Protocol and RDF Query Language)^85^ can be used to query the VO and retrieve vaccine-related information based on specific features via a SPARQL endpoint^86^.

Figure 5 presents three query examples: (a) Find *Brucella* vaccines using DL query; (b) Find influenza vaccines licensed for use in chickens using DL query; and (c) Find all vaccines against *Bordetella pertussis* defined in the OHDSI vocabularies, including their IDs assigned by different resources. Query (a) identified 84 specific *Brucella* vaccines. Query (b) revealed 18 vaccines that can be used to immunize chickens against the *influenza* virus. Query (c) found 462 vaccines that prevent *Pertussis* infection defined in the OHDSI vocabularies, including 17 vaccines in the CVX code set and 145 vaccines in the RxNorm. The queries and their results are available at the Zenodo repository^47^.

**Figure 5.**
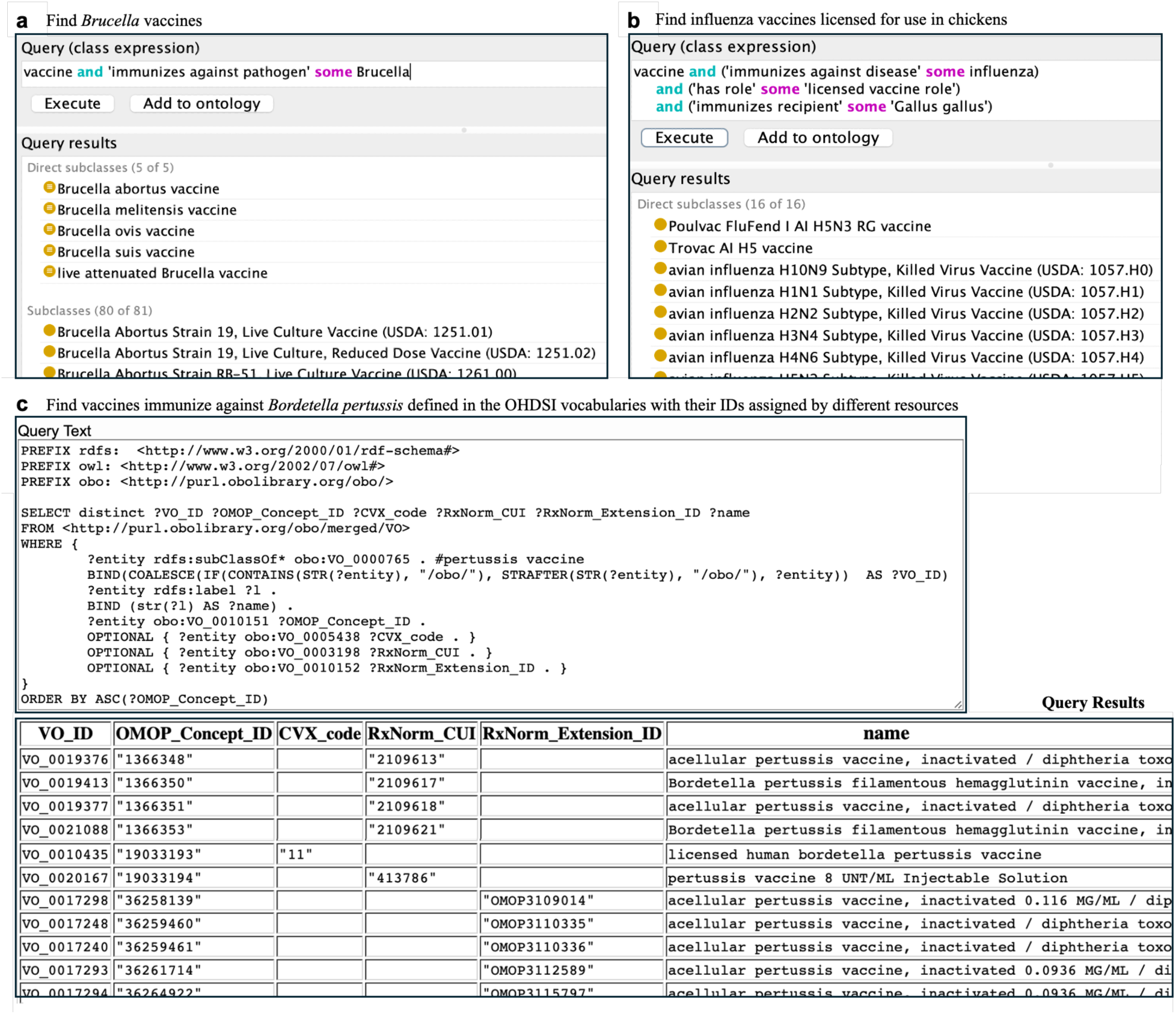
Semantic queries for vaccine identification based on vaccine features using DL and SPARQL query methods. (**a**) Find Brucella vaccines; (**b**) Find influenza vaccines licensed for use in chickens; (**c**) Find vaccines that immunize against *Bordetella pertussis* defined in the OHDSI vocabularies with their IDs assigned by different resources

The VIOLIN^55,56^ relational database contains a large amount of vaccine-related data. Our analysis found that the VO and VIOLIN represent overlapping information. For example, VO includes more licensed vaccine information imported from the RxNorm and RxE, while VIOLIN includes more detailed information about genes and proteins used for vaccine development and as vaccine immune factors^87^. VIOLIN also records detailed vaccine immune response studies from approximately 5000 journal articles or reliable websites, such as clinical trial studies from clinicaltrials.gov^10^. To integrate the complemented information from both resources, we have developed a vaccine knowledge graph (**VaxKG**) using Neo4j, which covers the information from both VO and VIOLIN^88^. Using Neo4j query language Cypher, we were able to query over VaxKG and obtain results that could not be obtained with either resource taken alone^88^. We are currently in the process of adding to VaxKG more information from further resources, such as PubMed literature or other ontologies, in order to expand its usability further.

***(3) Application of VO for enhanced experimental vaccine response studies***

As described earlier, VO has been used to model various vaccine responses and vaccine investigation studies^54,58,59^. The ontological vaccine investigation modeling has resulted in the identification of various variables in vaccine immune response studies^58^ and vaccine pathogen-challenge experiment studies^54,59^, leading to enhanced understanding of vaccine immune response^45^ and enhanced vaccine protection meta-analysis^59^.

VO has aggregated and represented extensive knowledge of vaccine-induced immune responses identified from the VaxImmutorDB vaccine immune factor database^87^. VaxImmutorDB has stored 1,741 vaccine immune factors from 13 host types (e.g., human, mouse, pig, and fish). These immune factors are inducible by 154 vaccines against 46 pathogens.

VO is the default ontology used by ImmPort in standardizing vaccine records^74^. As of March 3, 2025, the ImmPort repository contains 177 studies out of 1,188 investigating immune responses to vaccines or vaccine adjuvants. These recorded studies cover research on 69 vaccines and 13 adjuvants, and all of which are mapped to VO codes. With the VO identifiers of these vaccines, we were able to easily extract VO terms from the VO and generate a VO subset with a hierarchical structure (Figure 6) using the OntoFox tool^89^. The resultant VO-based classification allows us to gain a deeper understanding of the types of vaccines and adjuvants studied in ImmPort (seen in Figure 6a). Different types of COVID-19 vaccines, including mRNA COVID-19 vaccines, recombinant viral vector COVID-19 vaccines, and COVID-19 protein vaccines, were used in various ImmPort studies (Figure 6b), which enables us to explore immune responses biased to specific COVID-19 vaccine types. In addition, ImmPort records studies on different types of vaccine adjuvants (Figure 6c), facilitating vaccine adjuvant hierarchical analysis and deep mechanistic investigation. The VO terms used in ImmPort, the number of studies using each VO term, and the VO subset file are available at the Zenodo repository^47^.

**Figure 6.**
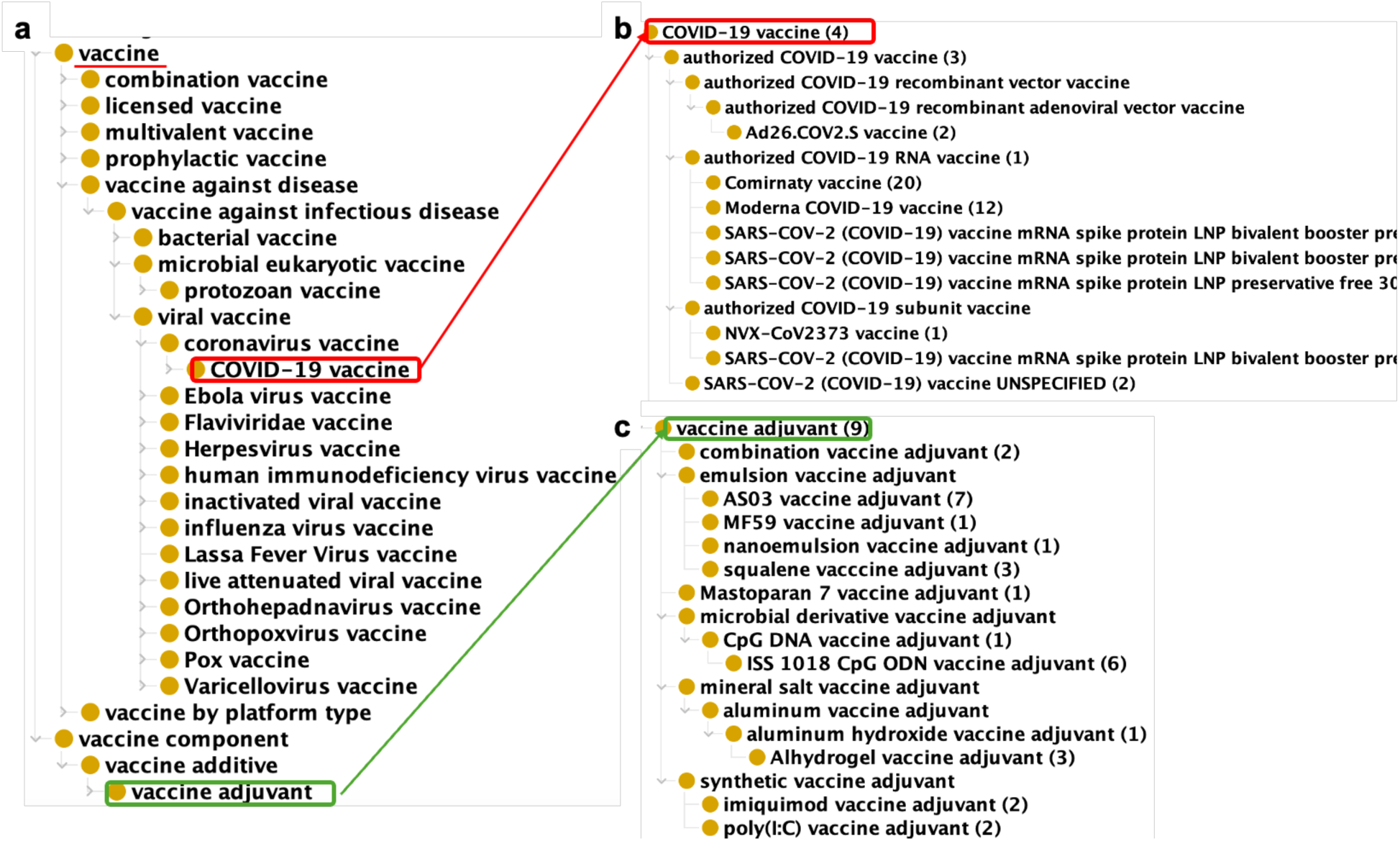
VO terms used in ImmPort. (**a**) VO hierarchy of vaccines and adjuvants in ImmPort. (**b**) COVID-19 vaccine terms with number of studies in ImmPort. (**c**) Vaccine adjuvants with number of studies in ImmPort.

VO has been used also in the development of VIGET^90^, a web portal for studying of vaccine-induced host responses based on Reactome pathways^91,92^ and ImmPort data^74^. VIGET uses the VO hierarchy to classify groups of vaccines. Since the raw omics gene expression data from ImmPort studies are stored in the GEO database rather than in ImmPort itself, we developed an ontology-annotated pipeline to automatically map ImmPort records to the raw gene expression data housed in the GEO^93^. This pipeline links the metadata represented in ImmPort with the gene expression data stored in GEO, enabling efficient data annotation and analysis. For example, we have used this pipeline to identify many vaccine-specific and sex-specific pathways from studies reported in ImmPort^90,93^.

***(4) VO-supported clinical vaccine data annotation and analysis***

VO has been used in tandem with the Ontology of Adverse Events (OAE)^94^ to support vaccine adverse event research. For example, by combining VO and OAE, we found distinct adverse event (AE) patterns associated with trivalent inactivated influenza vaccines (TIV) and live attenuated influenza vaccine (LAIV)^95^. Specifically, 13 out of 48 TIV-enriched AEs were shown to be closely related to specific neurological and muscular effects, including paralysis, movement disorders, and muscular weakness; and 15 out of 68 LAIV-enriched AEs were associated with inflammatory response and respiratory system disorders. Compared to TIV, LAIV was found to have a much lower chance of association with two severe adverse events – Guillain-Barre Syndrome and paralysis^95^.

The OHDSI initiative is a multi-stakeholder, interdisciplinary, open-science collaborative whose goal is to bring out the value of health data through large-scale analytics. Since January 2023 two key VO developers, Drs. He and Lin, have served as co-chairs of the OHDSI Vaccine Vocabulary Workgroup, whose overall goal is to utilize VO in enhancing the vaccine-related OHDSI data analysis. Unlike ImmPort, which directly uses VO for vaccine annotation, the OHDSI (OMOP) system uses RxNORM and RxE as standard vocabularies for vaccine terms.

However, RxNorm and RxE deploy a list structure, which means that the hierarchical features of vaccines which are captured in VO are left out of scope. OHDSI has also used the CDC CVX system for vaccine classification, but the results are then limited by the small number of CVX terms. VO, in contrast, includes all RxNorm and RxE vaccine terms and provides a solid hierarchical classification. VO also offers four unique vaccine classification options, namely: by targeted disease, by targeted pathogen, by mode of production, and by salient seasons of vaccination for influenza vaccines^66^. We have used the VO to study vaccine electronic health records (EHR) stored in OMOP-compliant resources, including the NIH National COVID Cohort Collaborative (N3C) data enclave, which is one of the largest clinical data resources for accelerating and collaborating on COVID-19 research^96^. For the latter, our study found that by using the VO hierarchy one can identify an increased number of reported vaccine EHR cases^97^.

Furthermore, VO can be used to study patterns in the data oriented around specific vaccine types such as mRNA vaccines or protein vaccines. Further collaboration with the OHDSI vocabulary team is ongoing to finalize the inclusion of VO in the OHDSI vocabulary system as a vaccine classification program^98^.

***(5) VO-based vaccine literature mining***

The VO has been used to greatly enhance vaccine-related literature mining. For example, the tool SciMiner supports literature indexing and gene name tagging using dictionary- and rule-based approaches^99^. Our study found that integrating VO significantly enhances SciMiner’s detection of vaccine-related literature and vaccine-gene interactions^100^ and vaccine-related gene-gene interactions^101^. As another example, a recent study found that a refinement using the VO enhanced the performance of ML algorithms in tagging vaccine names from clinical trial records^102^.

The primary reason for the enhanced performance of VO-base literature mining is its systematic hierarchical classification of vaccine types. For example, VO stores over 80 specific *Brucella* vaccines as subclasses of ‘*Brucella* vaccine’. *Brucella* is an intracellular bacterium that causes brucellosis, the most common zoonotic disease worldwide. In contrast, the Medical Subject Headings (MeSH)^103^, which is the vehicle for annotating PubMed documents, does indeed contain the term ‘*Brucella* vaccine’, but it includes zero subclasses under this term (https://meshb.nlm.nih.gov/record/ui?ui=D002004). This MeSH information offers limited support for searching for articles related to ‘*Brucella* vaccine’ in PubMed. As an illustration, our previous study showed that the search using this VO information dramatically increased the recall of searching “live attenuated *Brucella* vaccine” by 12-fold compared to search without using VO, while maintaining high precision (99%)^100^.

VO has also been used in combination with other ontologies to support advanced literature mining. For example, VO has been used together with the Interaction Network Ontology (INO) to support literature mining and analysis of vaccine-associated gene interaction networks^104^. Using 265,969 vaccine-associated documents from PubMed, 6,116 we identified gene pairs associated with at least one INO term, and 78 INO interaction terms were associated with at least five gene pairs. By comparing the background data set identified from 23,481,042 PubMed documents, 14 INO terms were significantly over-represented and 17 under-represented based on a modified Fisher’s exact test. Distinct vaccine-specific patterns were identified by analyzing these over- or under-represented INO terms under the INO hierarchy. For example, ‘nucleic acid cleavage’ and ‘glycosylation reaction’ were over-represented in vaccine-specific studies, and ‘phosphorylation’ and ‘dephosphorylation reaction’ were under-represented compared to the background signal. Such analyses provided insights for further vaccine-focused research.

The VO license allows for commercial adoption, widening the reach of the public ontology. SciBite from Elsevier^105^ offers the content of VO in multiple formats to create customer analytical solutions for different use cases. To start with, the full original source ontology is supplied in SciBite’s ontology management software, so that customers may either augment VO with custom terms as needed or use it in downstream applications such as electronic lab notebooks (ELNs), form filling and more. Additionally, SciBite provides a NER module based on the “vaccine adjuvant” branch of VO (VO:0000580). Adjuvants can modify the immunological response to a vaccine, thereby potentially decreasing the amount of antigen needed and/or the number of administrations necessary to achieve immunity. However, they may also trigger adverse events. Therefore, the study of their efficacy and safety is of special interest to biomedical and pharmaceutical communities. The SciBite vaccine adjuvant module was created in 2023 to fill an identified gap in the content that the company offered in that area, and is enriched with custom proprietary entities and synonyms. SciBite clients include large pharmaceutical and life-science companies, but also expand into non-biotech areas, and all benefit from the existence and development of a public resource such as the VO.

## Discussion

This report is the first manuscript to systematically introduce the development and applications of the Vaccine Ontology (VO), which has become the leading community-based vaccine standardization resource. VO integrates into its semantic framework vaccine related terms and associated definitions from multiple resources, including vocabularies, terminologies, literature, and websites. Many biological and biomedical resources and databases, such as VIOLIN^55,56^, ImmPort^74^, IEDB^76,77^, VAC^44^, and HIPC^75^, have used VO to harmonize vaccine related data. As a result, VO has made significant contributions to enhancing data FAIRness.

As a collaborative community-based effort, VO development has involved close collaboration with many other biomedical ontology initiatives. For example, the earlier stage of VO development involved close collaboration with the development of the Ontology for Biomedical Investigations (OBI)^15,16^. Several key terms such as ‘vaccine’ and ‘vaccination’ were defined in VO on the basis of close communication with the OBI team, and the early stage of OBI ‘assay’ modeling also utilized a vaccine investigation use case^15^. As another example, the VO team has worked with the Ontology for Parasite Lifecycle (OPL) development team by incorporating parasite lifecycle classes in VO’s account of vaccine development^70^. The combined application of VO and OPL provides a superior means for modeling and analyzing novel strategies for developing vaccines targeted to different stages of the life cycle of parasitic pathogens^70^.

Beyond the five areas of VO applications reported in this manuscript, VO has the potential to significantly advance our systematic understanding of vaccine mechanisms and to facilitate vaccine development. For example, we have started to categorize and define a number of different types of vaccine-induced immune responses and immune profiles by referencing newly identified scientific findings and wide discussions in the vaccine community^53^. In addition, our VIOLIN and VO projects have collected data on thousands of vaccine antigens^106^ and vaccine-induced immune factors^87^. Further ontology-based AI analysis of these data presents new opportunities to uncover how vaccines work and what types of immune pathways vaccines induce, which may suggest new directions for future research.

With the wealth of vaccine data accessible from the literature and other public resources, integrating all available vaccine data into the VO presents a significant challenge. To address this, we have developed templates and modules and generated a Makefile to automate importing terms from external resources as vehicles for creating new ontology terms in VO. Nevertheless, vaccine data collected from various sources, particularly from literature, still requires manual curation. This process is time-consuming and slows VO development, though it fosters high-quality standards in its results. Efforts to automate the integration of new vaccines into the VO are ongoing. Recently, we developed VaxLLMs, a tool that automates the annotation of *Brucella* vaccine data from literature^107^. This tool efficiently retrieves structured *Brucella* vaccine data from the literature with high performance. In the future, we plan to enhance the functionality of VaxLLMs and incorporate it into the broader VO development pipeline.

While large language modeling is a generative AI method currently being used extensively in research, large language models (LLMs) are only the beginning of the myriad possibilities that ML can contribute to advancement in modern scientific discovery. The drug R&D pipeline is a flow of research activities that travel from machine-assisted hypothesis generation with computational biology and laboratory experiments to compound validation and large-scale manufacturing and quality assurance and control. This process is also monitored by a robust regulatory framework. This complex environment of biomedical research activities calls for a hybrid approach requiring human expertise in diverse domains (including algorithm development, advanced statistical analysis, semantic technology, laboratory and in vivo experimentation) alongside computational infrastructure that can scale appropriately. To understand how best to construct this hybrid solution, one must map the various ML/AI methods to the different steps along this drug R&D pipeline. It will then be realized that this hybrid solution must be established on the solid foundation of well-understood data, primarily organized through ontological representation alongside predicted or reconstructed linked machine-readable graph representations for unstructured data. VO applications have already entered this next cycle of technology advancement through generative AI^88,102^.

Many directions exist for future VO development. Currently, VO primarily focuses on the representation of vaccines in English-speaking countries. A future effort is to extend VO’s inclusion of licensed vaccines from non-English speaking countries and regions. This inclusion will allow advanced integration and intelligent analysis of the large amount of vaccine data produced around the world. We are also developing LLMs to improve our literature mining performance in order to capture more data pertaining to vaccines information about which is recorded in the literature. We plan to standardize every vaccine record in clinical trial databases such as clinicaltrials.gov, but achieving such an ambitious goal will require significant effort from a broadened repertoire of stakeholders. Meanwhile, new applications can be developed to demonstrate further the power of VO in supporting advanced vaccine research and development in the exciting AI era.

## Methods

### Collaborative development to reach consensus in VO

In the early three years, domain and ontology experts met regularly in virtual meetings and annual face-to-face meetings were also held. During this period, key terms in vaccinology were identified and incorporated into the VO, and ontology design patterns were discussed and developed. Once the high-level hierarchy and design patterns were established, meetings were held to address emerging issues.

Beginning in January 2024, we held weekly meetings, focusing on reviewing the VO’s high-level classes and design patterns. These sessions improved class definitions and design patterns, ensuring consistent representation within VO.

Since January 2023, we have also organized regular bi-weekly meetings as part of the OHDSI Vaccine Vocabulary Workgroup. These meetings emphasize the hierarchical classification of licensed human vaccines using VO and their application in EHR data analysis^66,97,108^. We are closely collaborating with the OHDSI vocabulary workgroup, aiming to enhance and ultimately integrate the licensed VO section into the OHDSI vocabulary^98^.

VO developers and external users were encouraged to submit a term request or report issues through the VO issue tracker on GitHub (https://github.com/vaccineontology/VO/issues); this mechanism to request changes or additions has actively been maintained to this day.

### VO development strategies

VO is a community-based ontology in vaccinology that adheres to the OBO Foundry principles (e.g., openness and collaboration)^25^ and eXtensible Ontology Development (XOD) strategy^109^. The latter includes four basic methods: ontology term reuse, semantic alignment, use of ontology design patterns for new term generation, and community effort^109^.

**Supplemental Figure 1** presents the VO development framework. Once vaccine data are collected from various sources, they undergo standardization by mapping them to the VO and other OBO Foundry ontologies. If terms already exist in VO, no further action is needed. If terms are found in other OBO Foundry ontologies, they are imported into the VO using the OntoFox tool^89^. If any terms are missing from these ontologies, we create the needed terms and integrate them into VO by assigning VO IDs and aligning them with the VO hierarchy using the ROBOT tool^68^.

A combination of top-down and bottom-up methods was used in VO development. The top-down approach was initiated using the Basic Formal Ontology (BFO) as an upper-level ontology^29^, a framework widely adopted by the OBO Foundry and other ontologies to facilitate interoperability^24,28^. Domain experts identified the key terms in the field of vaccinology which were then defined and aligned with BFO and other OBO Foundry ontologies.

The bottom-up approach concerns vaccine data gathered from a wide range of sources to ensure comprehensive coverage, encompassing both human and nonhuman animal vaccines at various stages of development and approval. For human-licensed vaccines, data is sourced from established drug terminologies, including RxNorm^6^ for U.S.-approved vaccines and RxE^7^ for non-U.S.-approved vaccines. Additionally, vaccine information is sourced from the FDA vaccine site^5^ and the CDC CVX code set website^4^. The FDA approves vaccines that are licensed for use in the United States. CVX codes are part of the Health Level Seven International (HL7) standards and are used to standardize vaccine data across various healthcare systems. Licensed veterinary vaccines are retrieved from the USDA product catalog^9^, ensuring a thorough registry of vaccines used in nonhuman animals. Vaccines that are under development are primarily identified through scientific publications, providing insights into experimental and emerging vaccine technologies. Those in clinical trials are tracked using data from clinicaltrials.gov and NCI Thesaurus (https://ncithesaurus.nci.nih.gov/), which offers detailed information on cancer vaccine candidates being tested in various phases of clinical evaluation. Developed by the National Cancer Institute (NCI), the NCI Thesaurus^11^ is a comprehensive reference terminology for cancer-related research, and the vaccines identified within it are cancer vaccines.

This multi-faceted approach to vaccine data collection ensures that researchers and healthcare professionals have access to reliable and up-to-date information on vaccines across both human and veterinary domains. It encompasses vaccines at various stages of development and targets not only infectious diseases but also other conditions, such as cancer.

### Reuse of existing ontology resources

To establish an ontology interoperable with OBO Foundry ontologies, VO was developed using Basic Formal Ontology (BFO), a domain-independent ontology, as an upper-level ontology and reusing ontology terms from other existing ontologies. In addition VO utilized the OBO Metadata Ontology (OMO)^110^, an ontology of specified annotation properties that are used to annotate ontology terms for all OBO Foundry ontologies. The relation terms defined in the Relation Ontology (RO) were employed to represent commonly used relations^111^. RO Core is a small subset of RO containing the widely used relations in all OBO Foundry ontologies. The full versions of BFO, OMO, and RO core relations were imported into VO using OWL ontology importing. Other reused terms from existing ontologies were extracted and then imported on a term-by-term basis using OntoFox (http://ontofox.hegroup.org/)^89^.

### Adding to and editing of VO

Initially, VO terms were manually added to VO and edited using Protégé OWL editor (http://protege.stanford.edu/)^81^. However, adding many terms manually is time-consuming, error-prone, and requires OWL knowledge. To facilitate the efficient addition of terms to VO, ODPs were developed to create patterns for various VO terms^42,72^. The terms and their associated data that follow the same design pattern were formalized in a structured template in tabular format. The ROBOT tool template feature was used to convert the template into an OWL format file^68^. The terms were added or edited by making changes in the template and rerunning the ROBOT tool^68^. The Protégé editor 5.6.4 was primarily used to visualize and check the consistency of the ontology using ELK 0.6.0 reasoner^52^.

### Ontology release and quality control

The VO is released regularly, every 1-2 months. The latest VO OWL file is always available at http://purl.obolibrary.org/obo/vo.owl. The release pipeline was established and automated using a Makefile, which streamlines the workflow. The process begins by merging multiple OWL files into a single file. Next, a file with the fully inferred hierarchy is generated using the ELK 0.6.0 reasoner. The final released VO OWL file includes a specific URI, with the release date as the version IRI. This means specific releases remain available through this permanent, dated URL, and provides stability to users who can try newer releases and revert to a previously dated one if needed. As part of the release process, the ELK reasoner is utilized to verify ontology consistency and generate the inferred hierarchy, while the ROBOT tool’s report feature is employed to check the quality of the release file^68^, ensuring compliance with OBO Foundry principles.

Beyond the standard release quality checks conducted with ROBOT, an automated lexical-based tool^112^, is employed to identify missing “is-a” relations and duplicate terms, supporting further quality assurance. This tool operates by constructing “linked” and “unlinked” concept pairs based on shared lexical features, and generating Acquired Term Pairs (ATPs) to capture their unique word differences. When an ATP from a linked pair matches one from an unlinked pair, it suggests a potentially missing “is-a” relation. Such candidate relations undergo expert review and validation before incorporation into VO. This ATP-based tool is applied to the inferred hierarchy of the ontology, complementing the ELK reasoner.

### SPARQL and DL queries

The ontology was queried using both the Simple Protocol and Resource Description Framework (RDF) Query Language (SPARQL) and DL languages. SPARQL queries were executed via Ontobee’s SPARQL endpoint^86^, while DL queries were conducted using the DL Query tab in the Protégé OWL editor^46^.

### The VO applications

A systematical literature survey was performed to identify all the papers citing the VO. Representative VO applications were reported in five areas including harmonization/standardization of vaccine related data, vaccine knowledgebase development and queries, experimental vaccine study modeling and data analysis, clinical vaccine data annotation and analysis, and text mining.

## Data availability

The release version 2025-07-06 of VO presented in this paper is available at: http://purl.obolibrary.org/obo/vo/releases/2025-07-06/vo.owl. The latest version of VO is available at: http://purl.obolibrary.org/obo/vo.owl. In addition, VO is available on the Ontobee (http://www.ontobee.org/browser/index.php?o=VO), NCBO BioPortal (https://bioportal.bioontology.org/ontologies/VO), the Ontology Lookup Service (OLS) (https://www.ebi.ac.uk/ols4/ontologies/vo?tab=classes). The VO’s metadata is listed on the Bioregistry https://bioregistry.io/vo and on the OBO Foundry website https://obofoundry.org/ontology/vo.html. The VO is released under Creative Commons Attribution (CC BY) 4.0 License.

The mappings between VO and other vaccine related resources based on the SSSOM format are available on: https://github.com/vaccineontology/VO/tree/master/mappings.

The statistics, query results, and ImmPort use case described in the paper are all based on VO release version 2025-07-06^62^. The VO 2025-07-06 release file, along with source code, mappings files, statistic information, sample DL and SPARQL queries with their results, and ImmPort use case (including the VO subset file and a summary of VO annotations) presented in the paper, are freely accessible on the Zenodo repository (https://zenodo.org/records/16283855)^47^.

## Code availability

The VO code is available on the GitHub website: https://github.com/vaccineontology/VO. The VO code and use cases presented in this paper is also available on the Zenodo repository with the link provided above^47^.

The VO code is under Creative Commons Attribution (CC BY) 4.0 License.

## Author Contributions

JZ, AYL, AH, RR, KR, XingL, YP, ZX, and YH added terms to VO ontology. EO, JHu, EA, HK, XZ, NK, JG, VH, JS, AIS, SSay, FC, and XinnL collected vaccine information for VO addition. WWL and NK supported the representation of VAC vaccine adjuvants in VO. BP, AR, ADD, RHS, MC, PLW, MAM, CM, BDA, GSO, LGC, and BS supported early stage VO modeling. JZ, AH, AMM, BS, and YH finalized ontology design patterns. JZ, AYL, WM, RA, YP, KH, AH, AD, QY, JHur, and YH participated in collaborative OHDSI vaccine vocabulary workgroup study. WM, RA, and LC supported ontology term mapping and quality assessment. RV, CTH, and PR requested new terms and identified VO issues. JZ and CTH prepared SSSOM format mapping files. AH, GW, SSarn, and TT, FYY supported VO-based data analysis. JL, HR, BB, RPH, AO, CT, and JHur supported VO-based literature mining. BS ensures VO is aligned well with the Basic Formal Ontology. AMM, BP, LAC, HLTM, GSO, LGC, and YH served as vaccine domain experts. YH initiated and led the VO development. JZ and YH prepared the first draft of the manuscript. YH prepared Figure 2, and JZ prepared the other figures, Table 1, and supplemental materials. All authors participated in paper editing and discussion. All authors reviewed the manuscript and agreed on its publication.

## Acknowledgements

This research is supported by NIH grants U24AI171008 (YH, JHur, and CT), R01AI081062 (YH), UH2AI132931 (YH and GW), P30ES017885-01A1 (GSO), U24CA271037 (GSO), U01AI167892 (BP and RV), HHSN266200400041C (RHS), N01AI2008038 (RHS), N01AI40076 (RHS), R01LM013335 (LC), R01NS116287 (LC), U24AI177622 (LGC), by a Rackham Pilot Research project at the University of Michigan (YH), by the National Science Foundation (NSF) through grant 2047001 (LC), and by the Public Health Agency of Canada / Canadian Institutes of Health Research Influenza Research Network (PCIRN) (MC). Strong support from the NCBO, NCIBI, the OBI Consortium, OBO Foundry, and IDO Initiative is greatly acknowledged.

## Disclaimer

RR currently works at the US Food and Drug Administration but participated in this effort during her time at the University of Michigan. This article reflects the views of the authors and should not be construed to represent either the FDA’s or the NIH’s views or policies.

