## Supplemental Figure 1 for "VO: The Vaccine Ontology"

### Supplemental Materials:

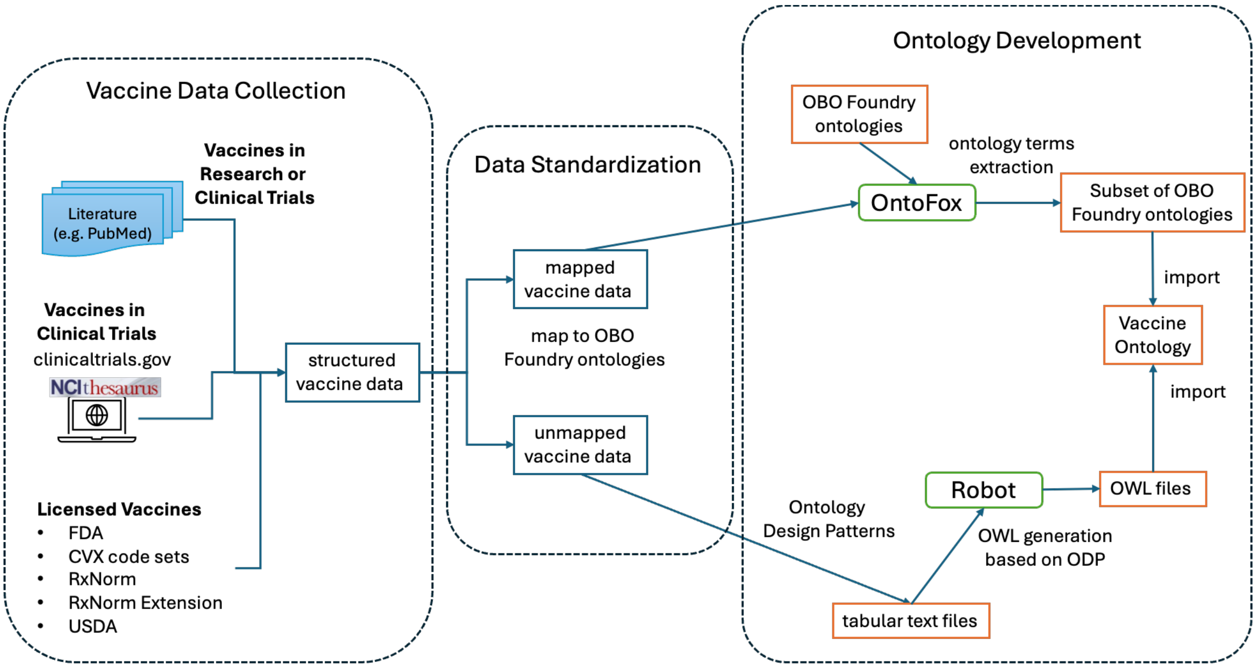


**Supplemental Figure 1**. Overview of VO development framework. Abbreviations: FDA: Food and Drug Administration; CVX: Centers for Disease Control and Prevention; USDA: U.S. Department of Agriculture; OBO: Open Biological and Biomedical Ontology; ODP: Ontology Design Pattern.
